# utR.annotation: a tool for annotating genomic variants that could influence post-transcriptional regulation

**DOI:** 10.1101/2021.06.23.449510

**Authors:** Yating Liu, Joseph D. Dougherty

**Affiliations:** Department of Genetics, Washington University School of Medicine, St. Louis, MO, USA; Department of Psychiatry, Washington University School of Medicine, St. Louis, MO, USA

## Abstract

**Summary:** Whole genome sequencing of patient populations is identifying thousands of new variants in UnTranslated Regions(UTRs). While the consequences of UTR mutations are not as easily predicted from primary sequence as coding mutations are, there are some known features of UTRs modulate their function. utR.annotation is an R package that can be used to annotate potential deleterious variants in the UTR regions for both human and mouse species. Given a CSV or VCF format variant file, utR.annotation provides information of each variant on whether and how it alters known translational regulators including:upstream Open Reading Frames (uORFs), upstream Kozak sequences, polyA signals, the Kozak sequence at the annotated translation initiation site, start codon, and stop codon, conservation scores in the variant position, and whether and how it changes ribosome loading based on a model from empirical data.

**Availability and implementation:** utR.annotation is freely available on Bitbucket (https://bitbucket.org/jdlabteam/utr.annotation/src/master/) and CRAN (to be updated)

**Supplementary information:** Supplementary data are available at https://wustl.box.com/s/yye99bryfin89nav45gv91l5k35fxo7z

## 1. Introduction

Post-transcriptional regulation is essential to the control of translation, controlling the abundance, timing, and location of new protein production. For example, key regulatory elements in untranslated regions (UTRs), such as upstream open reading frames (uORFs) (Barbosa et al., 2013; Calvo et al., 2009), Kozak sequences (Acevedo et al., 2018; Kozak, 1986), and polyA signals (Rehfeld et al., 2013; Weill et al., 2012) strongly influence protein production. Variants that alter these regulatory elements could substantially modify the final protein levels and potentially lead to disease. Thus, annotation of genomic variants in UTRs is essential to reveal potential etiopathogenetic mechanisms. Given the tremendous number of sequence variants being discovered, highlighting putative deleterious variants would be useful to prioritize functional studies, or weight human genetic analyses. Combined Annotation-Dependent Depletion (CADD) (Rentzsch et al., 2019) is widely used to measure variant deleteriousness, however its algorithm is currently unaware of many key regulatory features of UTRs. A recently developed UTR annotator(Zhang et al., 2020) only annotates uORFs perturbation in 5’ UTR. Therefore, we developed a R package, utR.annotation, that annotates genetic variants’ effect on additional categories of UTR key regulatory elements in both 5’ and 3’ UTRs.

## 2. A R package for annotating potential deleterious variants in non-coding regions

The utR.annotation package can be used to highlight potentially deleterious variants by annotating its impact on known UTR regulators easily predicted from sequence: uORFs, upstream Kozak sequences (uKozaks), polyA signals, the Kozak sequence at the annotated translation initiation site (aKozaks), as well as start codon, and stop codon. In addition, as highly conserved regions tend to have important biological functions, the package also provides Phastcons and PhyloP conservation scores at the variant positions (Fig 1A). Finally, the utR.annotation package also leverages a deep learning model trained from empirical data (Sample et al., 2019) to predict changes in mean ribosome loading based on 5’ UTR on reference and alternative alleles.

**Figure 1:**
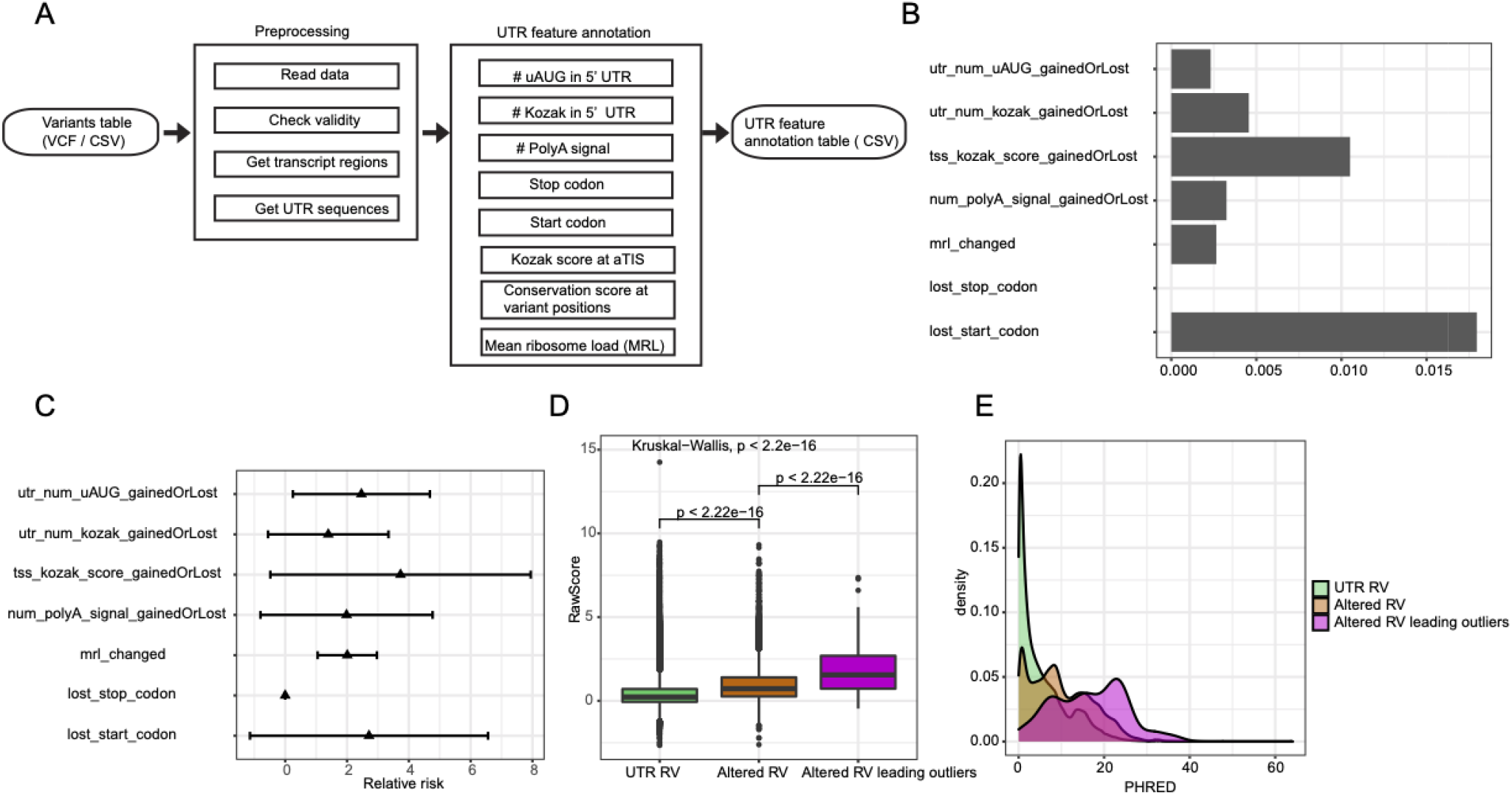
utR.annotation pipeline and its application on identifying high risk variants of eOutliers in GTEx. A) utR.annotation pipeline. B) Proportion of utR.annotation identified variants that were carried by eOutliers individuals. C) Relative risk of seven variant types. D) CADD raw scores comparison among three groups of variants. E) PHRED distributions of three variant groups.

### 2.1 Input and output formats

runUTRAnnotation function is used to annotate any genomic variant for potential UTR regulatory impact. As input, the user provides variant information in either the standard VCF or CSV format including four columns: Chr, Pos, Ref, and Alt. They also specify species, a reference genome build, and output directory. To increase speed, the user may optionally include a Transcript column as well, providing an ENSEMBL format transcript ID overlapping each variant, if already available. The annotation table is outputed in CSV format. The sample test data with input and output are provided in the supplementary data. Detailed explanation on each column of the output file is at CRAN & Bitbucket.

### 2.2 Robust on genome annotations

The utR.annotation works robustly on mouse and human variants regardless of genome build. runUTRAnnotation will use the latest Ensembl genome annotation by default and you can choose any available Ensembl version for identifying gene/transcript sequences.

### 2.3 Efficient on large data

For less than 10,000 variants, running the annotation on a typical PC is feasible and benchmarks at < 210 minutes with 1 CPU. For larger datasets, the tool can run in parallel on multiple cores (e.g., cluster) (Figure S1). Besides leveraging internal parallelism, a large variant file can be partitioned into a user specified number of small files using the partitionVariantFile function, and partitioned outputs can be concatenated with the concatenateAnnotationResult function. Using these tools, the GTEx example below (~23 million variants) took 54 hours when partitioned into 5000 small files and using 4 CPUs for each file.

## 3. Application on identifying high risk variants of eOutliers in GTEx

Rare variants with large effects tend to contribute to complex disease risk (Gibson, 2012). However, Identifying functional rare variants (RV) in the noncoding regions remains challenging. Many noncoding variant mutations are thought to influence gene expression, and UTR mutations in particular may influence transcript stability either directly (e.g., impacting miRNA binding sites) or indirectly (e.g., influencing the transcript initiation, as translation can either increase or decrease transcript stability). Either circumstance suggests the hypothesis that rare UTR mutations could contribute to unusual transcript levels in a given individual. Therefore, we annotated UTR rare variants (allele frequency < 1%) (uRV), and to identify the frequency with which these were associated with outliers in expression across a population of individuals (eOutliers) (Ferraro et al., 2020). The data set comprised 20,252 genes from 838 samples of transcriptome data in the Genotype-Tissue Expression (GTEx) project version 8 (v8). We used utR.annotation to annotate all RV from GTEx on genes in the eOutlier data and identified 59,986 uRV altering UTR regulatory elements (num_polyA_signal_gainedOrLost: 6,470, utr_num_uAUG_gainedOrLost: 7,782, utr_num_kozak_gainedOrLost: 1,536, tss_kozak_score_gainedOrLost: 1,904, lost_start_codon: 724, lost_stop_codon:414, mrl_changed: 41,156).

We next examined the proportion of these that corresponded to an unusual RNA level. We found that more than 1% of lost start codon variants (13/724=0.018) and aKozak strength changing variants (20/1904=0.011) were carried in eOutliers (Fig 1B), proportions that are higher than expected by chance. Overall aKozak strength changing variants had the largest relative risk (3.73 ± 4.21). All variant types except for lost stop codon variants had relative risk > 1 and were enriched in eOutliers (Fig 1C).(Note – no lost stop codon variant was found in eOutliers).

We next compared our annotation to CADD, a standard variant annotation tool. CADD combines numerous different variant features into a single score. It is aware of conservation scores in UTR features, but not ORF, Kozak, PolyA, and MRL. We compared the CADD scores of three groups of RVs: uRVs, uRVs that altered UTR feature elements, and uRVs that altered UTR feature elements and were carried in eOutliers. uRVs leading eOutliers have significantly higher CADD scores than the other two groups, indicating their higher deleteriousness (Fig 1D). Specifically, forty-five unique uRVs carried in eOutliers had PHRED scores larger than 20, which indicated they’re ranked as 1% most deleterious variants in humans (Table S1), suggesting some UTR information is being annotated by CADD. However, Our annotation also captured 55 potential deleterious variants that were carried in eOutliers but had CADD scores less than 10 (Table S2), suggesting CADD is missing some information. Four of the CADD underestimated variants altered ployA signals in 3’ UTR and 2 out of 4 have moderate conservation (phastCons = 0.653 and 0.351). One CADD underestimated variant changed MRL and was in a highly conserved position (phastCons = 0.994). One CADD underestimated variant altered uORF and has high conservation score (phastcons = 1). Overall, our utR.annotation could detect additional presumably harmful variants that are overlooked by current annotation tools, and thus might make a useful addition to the next version of CADD.

Finally, we’d note that there are many mechanisms by which a variant might be highly detrimental to protein production, but not influence the transcript levels (the only readout available currently for GTEx). Thus, this annotation tool may show even higher overlap in future studies of protein level outliers as well.

## 4. Conclusion

Our utR.annotation is a user-friendly and scalable tool to annotate variants that alter the UTR regulatory elements. It can be used to select top deleterious variants for functional analysis and provide detailed information on UTR interruption for further exploration, or statistical analysis.

## Acknowledgements

We would like to thank Paul Sample and the Seelig lab for providing a Kozak PWM and the MRL algorithm, and the Battle lab for the eOutlier lists. We would also like to thank Sergej Djuranovic, Tomas Lagunas, Tony Fischer, Stephen Plassmeyer, Stephan Sanders, and Michael Wells for helpful comments and discussion. This work was supported by the Simons Foundation (571009) and the NIH (5R01MH116999, 5R01NS10227, R01GM112824).

## Supplementary information

**Figure S1:**
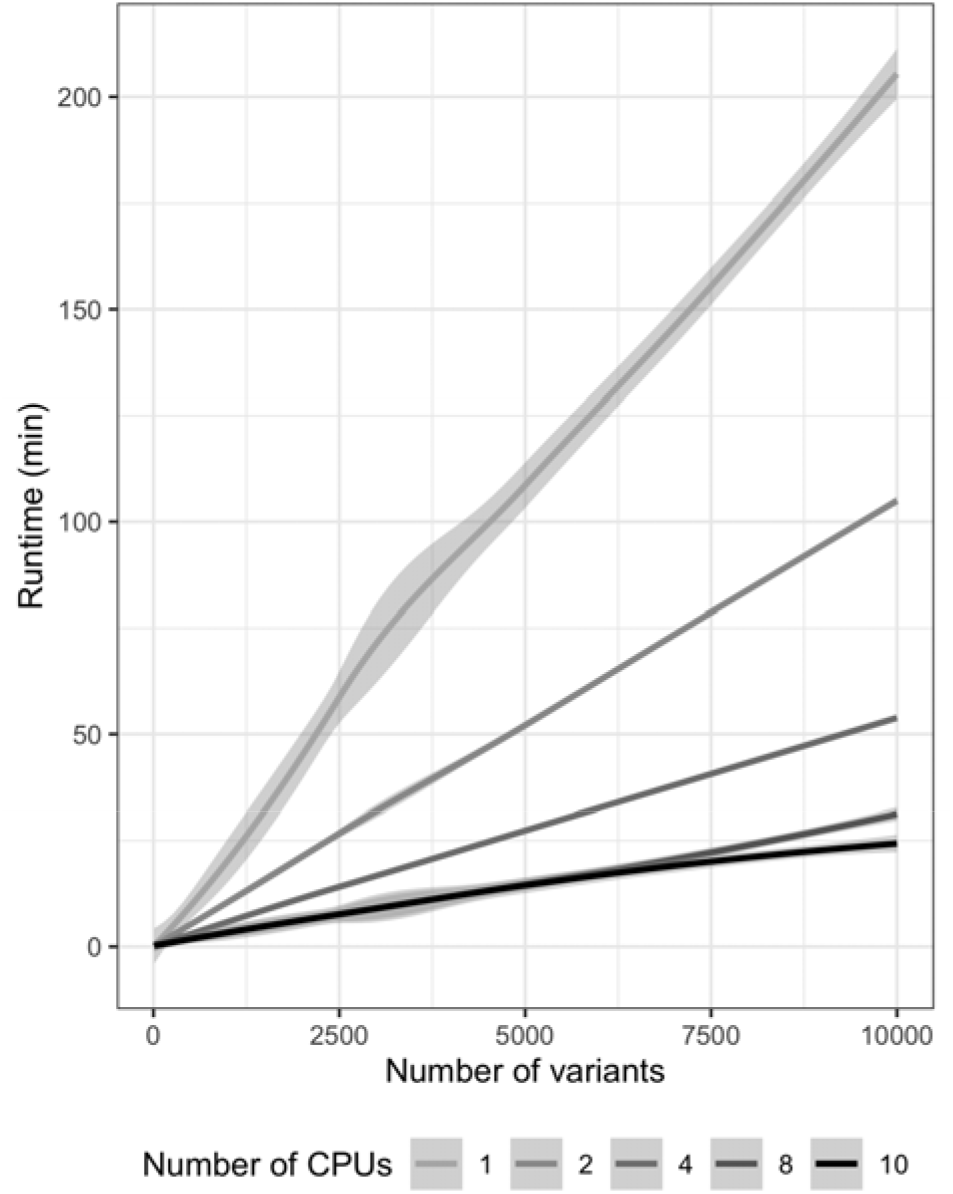
Runtime (min) of runUTRAnnotation on 10,000 variants with 1, 2, 4, 8 and 10 number of CPUs.

**Figure S2:**
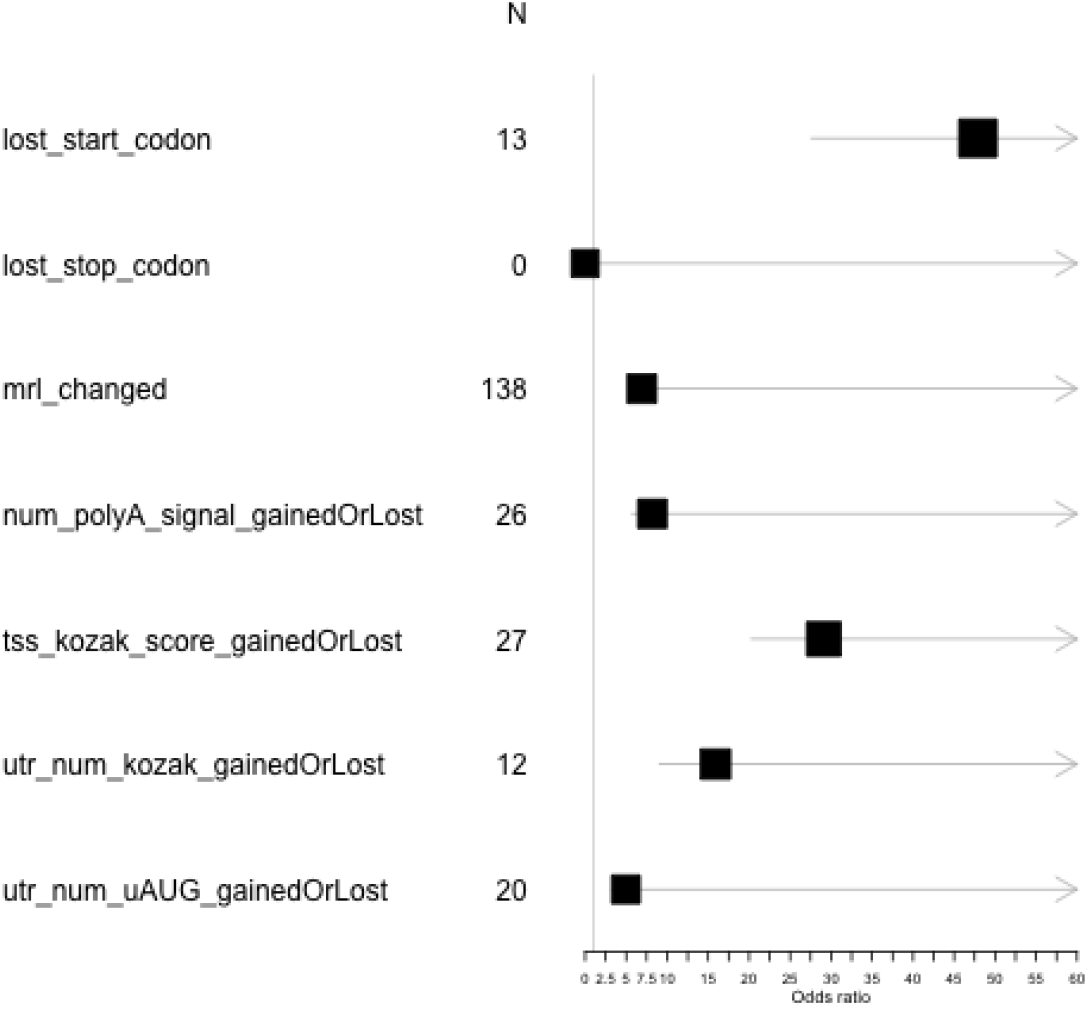
Fisher’s exact test on the association of the variant types and eOutliers

**Table S1: Top deleterious variants annotated by CADD**

Forty-five uRVs that carried in eOutliers and had PHRED scores larger than 20. The first 8 columns are chromosome, position, reference sequence, alternative sequence, CADD RawScore, CADD PHRED score, altered feature element, and RV group. The rest of the columns are annotation output from utR.annotation. The detailed description of those columns can be found on BitBucket.

**Table S2: Potential deleterious variants annotated by utR.annotation but missed by CADD**

Fifty-five potential deleterious variants that were carried in eOutliers but had CADD scores less than 10. The first 8 columns are chromosome, position, reference sequence, alternative sequence, CADD RawScore, CADD PHRED score, altered feature element, and RV group. The rest of the columns are annotation output from utR.annotation. The detailed description of those columns can be found on BitBucket.

## Methods

### Analysis of eOutlier data from GTEX

Expression outlier calling was performed by individual’s median Z scores of 785 individuals on 20,252 genes. The expression outliers (eOutliers) for each gene were selected with a threshold as | median Z score | > 3. We retained RVs from GTEx v8 VCF that passed quality control, had AF < 1% and located within any gene in the expression outliers data.

To assess how RVs in noncoding regions contribute to expression outlier patterns, we used utR.annotation package to annotate RVs for their impact on UTR regulatory elements using Ensembl v100 human annotation. The proportions of variants that were carried in eOutliers for six variant types were calculated and compared in Fig 1B. The relative risk of the variant types were compared in Fig 1C. We calculated the relative risk as the proportion of outliers with a given variant type over the proportion of non-outliers with the given variant type. The association of the variant types and eOutliers were shown in Fig S2. N represents the number of variants in the specified variant type that were carried in eOutliers. Odds ratios and confidence intervals were calculated by the Fisher’s exact test.

CADD scores (Genome build GRCh38 / hg38) were retrieved from all gnomad release 3.0 SNV / InDels downloaded from the CADD website and the CADD scores for remaining variants not found in gnomad data were queried using the CADD web server. We compared the CADD raw scores among three groups of variants in Fig 1D: 1) variants that are in UTRs; 2) variants that are in UTRs and annotated as one or more types of variants; 3) variants that belong to one or more variant types and also are carried in outliers. Kruskal-Wallis test and Wilcox test were performed for all groups comparison and pairwise comparisons, respectively. The distribution of CADD PHRED scores of the three groups were shown in Fig 1E.

## References

1. Acevedo JM, Hoermann B, Schlimbach T, Teleman AA (2018) Changes in global translation elongation or initiation rates shape the proteome via the Kozak sequence. Sci Rep 8:4018.

2. Barbosa C, Peixeiro I, Romão L (2013) Gene Expression Regulation by Upstream Open Reading Frames and Human Disease. PLOS Genet 9:e1003529.

3. Calvo SE, Pagliarini DJ, Mootha VK (2009) Upstream open reading frames cause widespread reduction of protein expression and are polymorphic among humans. Proc Natl Acad Sci 106:7507–7512.

4. Ferraro NM et al. (2020) Transcriptomic signatures across human tissues identify functional rare genetic variation. Science 369.

5. Gibson G (2012) Rare and common variants: twenty arguments. Nat Rev Genet 13:135–145.

6. Kozak M (1986) Point mutations define a sequence flanking the AUG initiator codon that modulates translation by eukaryotic ribosomes. Cell 44:283–292.

7. Rehfeld A, Plass M, Krogh A, Friis-Hansen L (2013) Alterations in Polyadenylation and Its Implications for Endocrine Disease. Front Endocrinol 4.

8. Rentzsch P, Witten D, Cooper GM, Shendure J, Kircher M (2019) CADD: predicting the deleteriousness of variants throughout the human genome. Nucleic Acids Res 47:D886–D894.

9. Sample PJ, Wang B, Reid DW, Presnyak V, McFadyen IJ, Morris DR, Seelig G (2019) Human 5′ UTR design and variant effect prediction from a massively parallel translation assay. Nat Biotechnol 37:803–809.

10. Weill L, Belloc E, Bava F-A, Méndez R (2012) Translational control by changes in poly(A) tail length: recycling mRNAs. Nat Struct Mol Biol 19:577–585.

11. Zhang X, Wakeling M, Ware J, Whiffin N (2020) Annotating high-impact 5′untranslated region variants with the UTRannotator. Bioinformatics.

