## Supplementary figures and images for "utR.annotation: a tool for annotating genomic variants that could influence post-transcriptional regulation"

### Figure S1

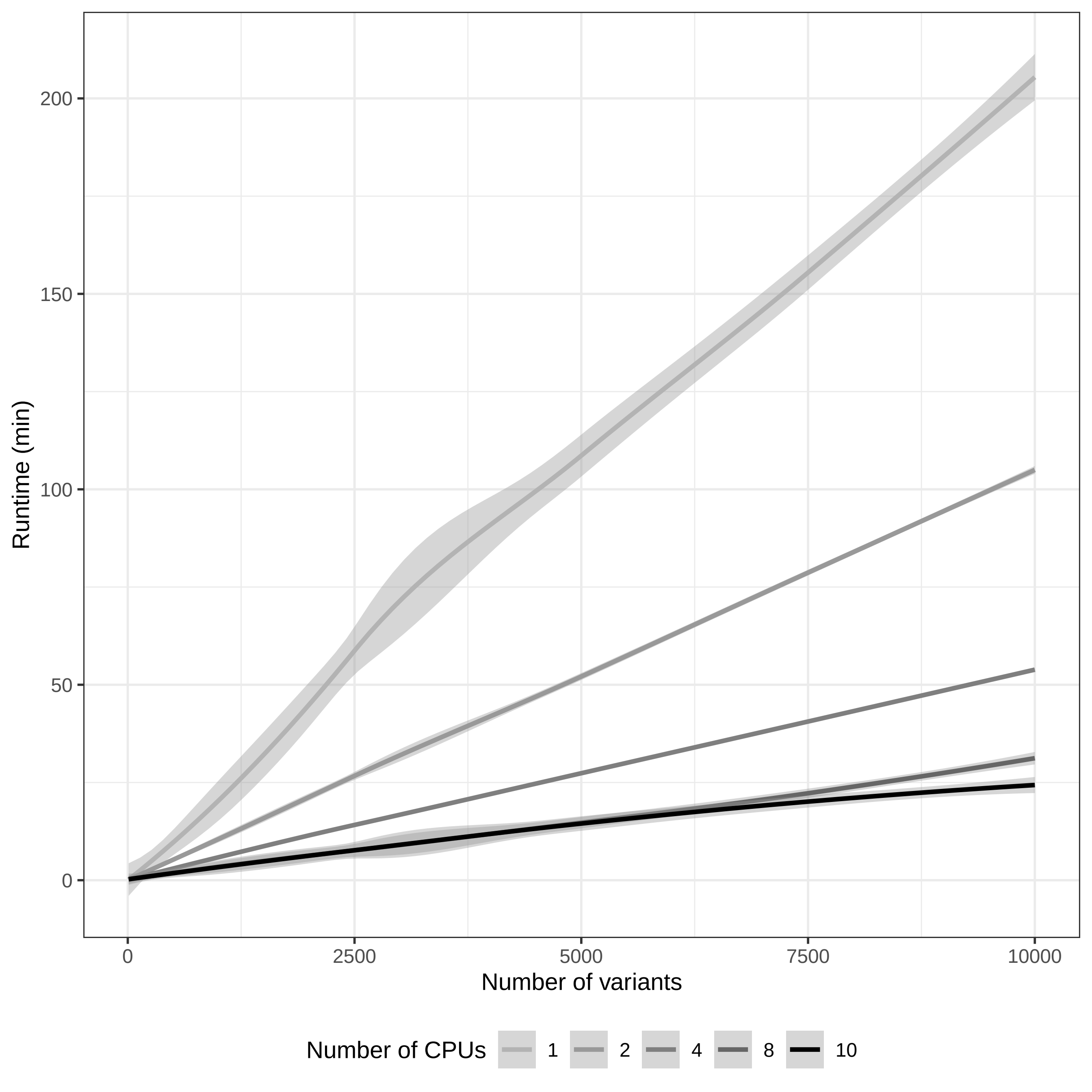

### Figure S2

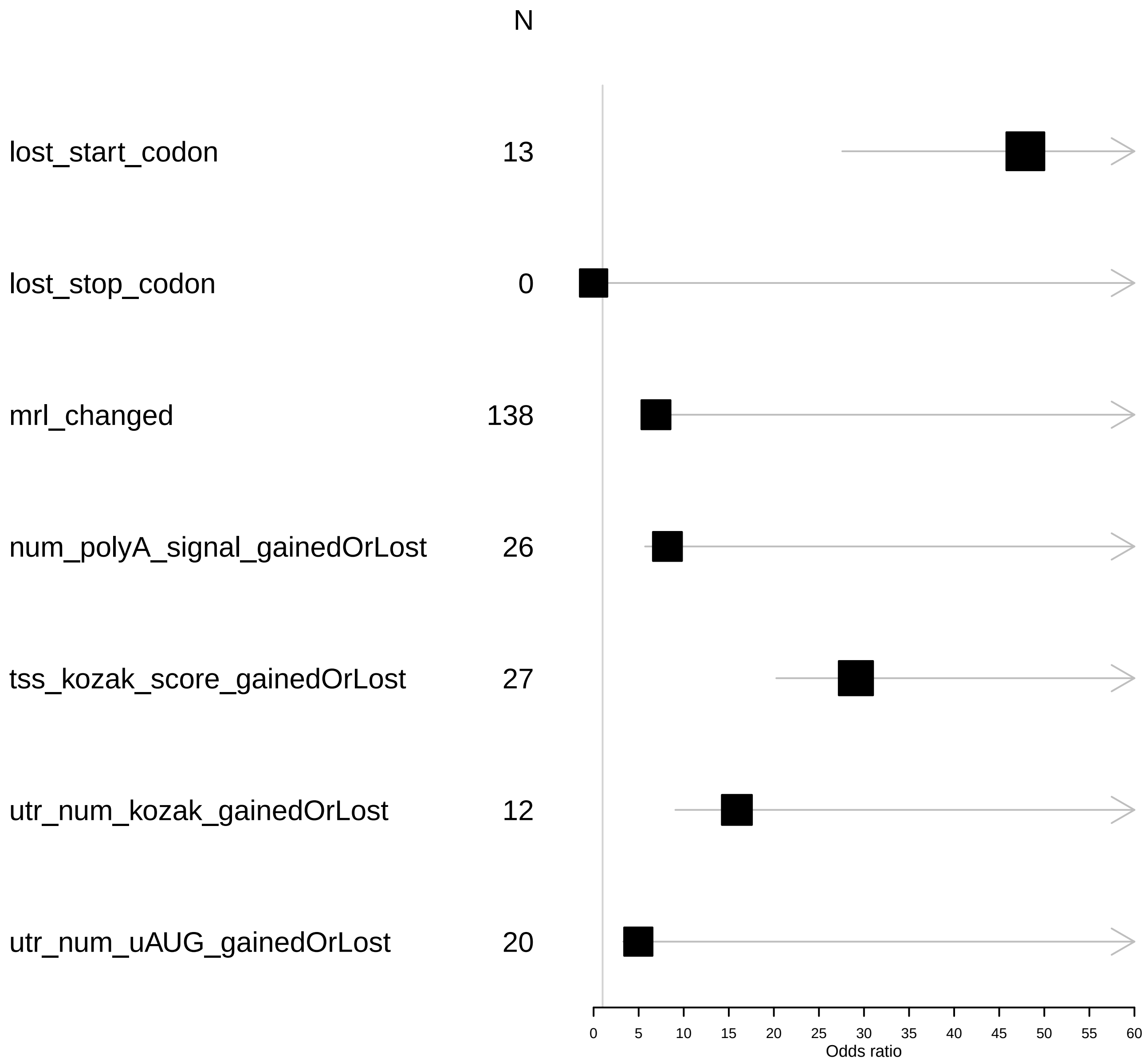
